# CalR: A Web-based Analysis Tool for Indirect Calorimetry Experiments

**DOI:** 10.1101/213967

**Authors:** Amir I. Mina, Raymond A. LeClair, Katherine B. LeClair, David E. Cohen, Louise Lantier, Alexander S. Banks

## Abstract

We report a web-based tool for analysis of indirect calorimetry experiments which measure physiological energy balance. *CalR* easily imports raw data files, generates plots, and determines the most appropriate statistical tests for interpretation. Analysis with the general linear model (which includes ANOVA and ANCOVA) allows for flexibility to interpret experiments of obesity and thermogenesis. Users may also produce standardized output files of an experiment which can be shared and subsequently re-evaluated using *CalR*. This framework will provide the transparency necessary to enhance consistency and reproducibility in experiments of energy expenditure. *CalR* analysis software will greatly increase the speed and efficiency with which metabolic experiments can be organized, analyzed according to accepted norms, and reproduced—and will likely become a standard tool for the field. *CalR* is accessible at *https://CalR.bwh.harvard.edu.*

**Graphical Abstract:** 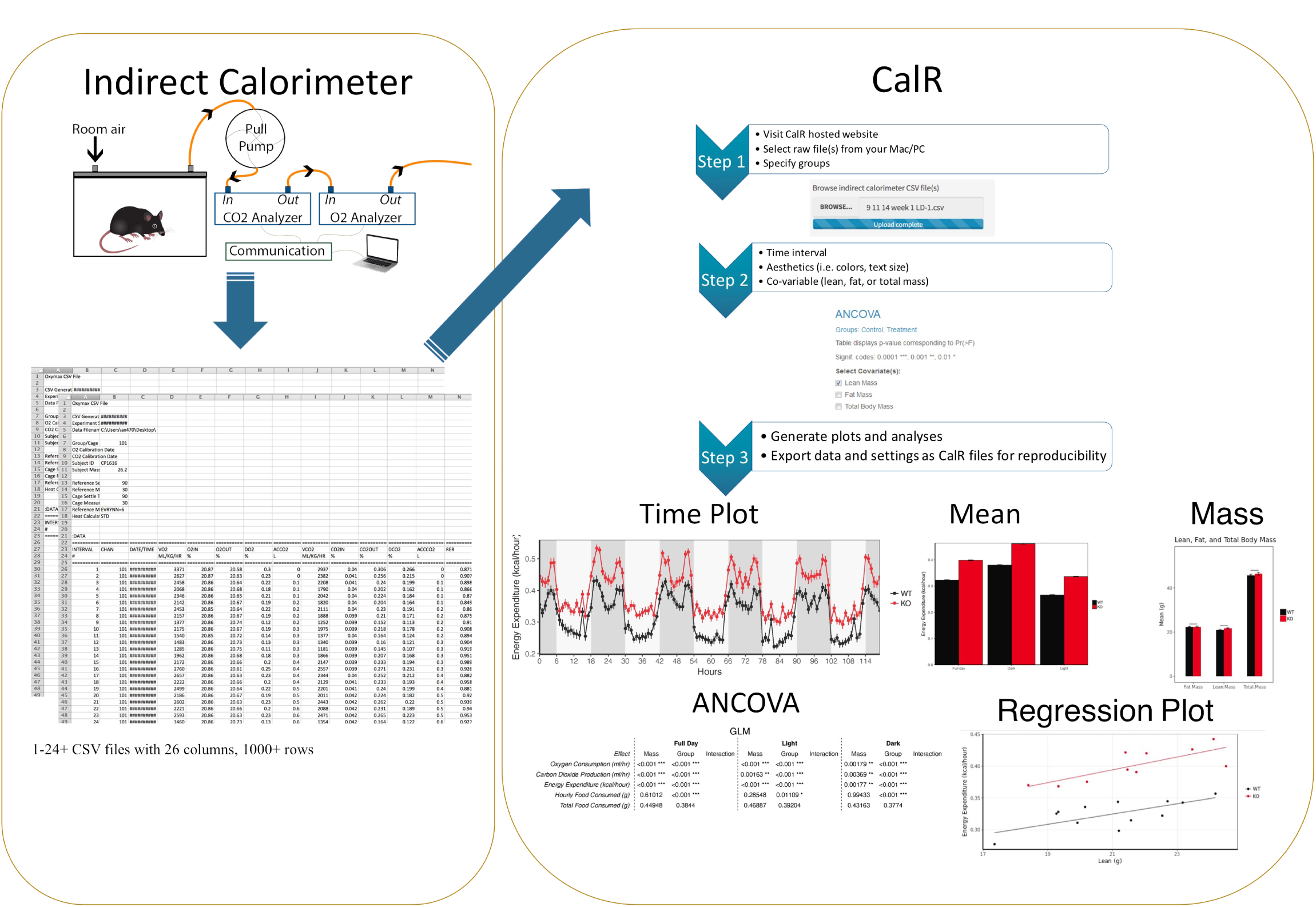

## Introduction

The increased prevalence of obesity, which arises with an imbalance in food intake and energy expenditure (EE), is driving increased morbidity and mortality worldwide [1]. While much focus has been placed on elevated energy intake as the primary force driving the rise in obesity, increasing attention is being directed toward the therapeutic potential to increase EE [2, 3]. In addition, decreased EE following weight loss contributes to the persistence of obesity [4]. Indirect calorimetry measurements of EE have proven invaluable in furthering our understanding of the pathogenesis of obesity [5]. Indirect calorimetry is not invasive compared to other methods of EE determination. Alternatives, such as direct calorimetry or doubly-labeled water analysis require sacrificing the animals and harvesting organs eliminating the possibility for serial measurements. In contrast, indirect calorimetry allows for more flexible and sophisticated experiments that can be repeated in the same animals over time. Although the use of indirect calorimetry has become widespread, controversies have emerged on the appropriate treatment of the data generated by these experiments, fundamentally challenging some published conclusions [6–8]. Because analysis of these large data sets is somewhat onerous, a need exists for a tool to assist with appropriate analysis and interpretation of results. The absence of such a tool has led to conflicting interpretations of experimental data [7, 8].

Following decades of debate, analysis of covariance (ANCOVA) has become the consensus approach to the treatment of indirect calorimetry EE data when comparing animals of different body composition as occurs in obesity [5, 9–11]. ANCOVA is a type of general linear model which uses both regression and ANOVA (Analysis of Variance) to adjust for covariates. Because neither lean body mass nor fat mass is metabolically inert, ANCOVA includes either LBM or FM as a covariate in the analysis of EE. However, in many cases, restrictions exist for widespread implementation of ANCOVA. These barriers include “wrangling” large data sets to prepare the raw data for analysis, unfamiliarity with statistical software packages, and the lack of a commercial software package to perform statistical analysis of indirect calorimetry experiments, despite the need for a solution long being apparent [5, 9–18]. As a consequence regression-based analysis, such as ANCOVA, is not consistently being implemented in the analysis of energy balance in mice of different body composition [6, 19].

While ANCOVA brings many benefits to the interpretation of energy balance, its use can be overly restrictive in experiments of non-shivering thermogenesis. Activation of Brown Adipose Tissue (BAT) increases EE through heat generation which is strongly dependent on BAT and lean body mass. The interaction between EE and lean body mass in experiments of thermogenesis may violate assumptions of the ANCOVA. We find the general linear model (GLM) allows for regression-based analysis of energy expenditure in experiments of obesity and thermogenesis (i.e. brown-fat mediated thermogenesis)[10].

Here we describe the open-source *CalR* software project, our effort to reduce the burden to perform statistical analysis of indirect calorimetry data by creating an easy to use software tool for the scientific community. With *CalR*, users can import large data files, evaluate experimental validity, examine data for experimental outliers, and compare differences between groups using GLM. The results are exportable as files that can be shared in a centralized repository or as supplementary data accompanying publications. In this manuscript, we focus on describing the application of this software tool to real-world examples for appropriate data analysis. *CalR* has the potential to become the standard resource for examination of energy balance experiments in laboratory animals.

## Methods

### Overview of the software system

This software package, designated *CalR (*an abbreviated form of *calor*, the Latin word for heat), is written in the R programming language using a *Shiny* graphical user interface (GUI) to capitalize on robust statistical analysis routines, free availability, and intuitive user interface. In partnership with Partners Healthcare, we hosted *CalR* as a publically available web application. Each analysis template is set up as a distinct Shiny application individually hosted on the server and routed to a web page within https://CalR.bwh.harvard.edu/.

### Software Architecture and Workflow

The primary functions of *CalR* broadly include reading and visualizing raw calorimetry data and performing statistical analysis. The user-friendly *CalR* web pages allow the user to specify body mass data and assigning subjects into groups. Navigating through the tabs of *CalR*, users will find their data for metabolic variables plotted either as group averages or as individual tracings. Subsequently, *CalR* conducts the appropriate data analyses depending on the metabolic parameter and using mass as a covariate as necessary. The abundance of input options gives users the flexibility to explore their data from a variety of experimental designs.

### System compatibility

The data generated by any of the three high-quality manufacturers of indirect calorimeter systems for small animals (Sable Systems, TSE, and Columbus Instruments) can be imported directly using *CalR*’s GUI.[20]. The “Input” tab in *CalR* contains a section in which a user may import one or more Comma Separated Value (CSV) files; this will depend on the manufacturer’ system. Below are specific steps for selecting the preferences to allow data import into *CalR* from each of these systems.

### Instructions for data preparation

#### Columbus Instruments’ CLAMS (Comprehensive Lab Animal Monitoring System)

A critical shortcoming of this system software is the “automatic normalization” which divides metabolic parameters by body weight (e.g., VO_2_ ml/kg/hour). When analyzing CLAMS data, *CalR* will reverse this normalization (e.g., VO_2_ ml/hour) before any further calculations. For this reason, when setting up the experiment by navigating to **Experiment > Setup**, users may enter any value for the subject mass and maintain the default “Volume Rate Units” setting to “ml/kg/hr” under **Experiment > Properties**. Here the user should also make sure the “Heat Calculation” setting is “Standard, kcal”. After an experiment has been completed and stopped, open Oxymax and select “Run Oxymax as Data Viewer”. When prompted, choose the hardware configuration file (.ini) used for setting up your experiment. Next, navigate to **File > Open experiment data** and open the .CDTA file from the CLAMS run. Once opened, navigate to **File > Export > Export all subject CSV’s**. Each cage run by the CLAMS system generates a separate output file.

#### Sable Systems’ Promethion

The high data density collected by Promethion systems necessitates pre-processing steps to reduce file sizes and processing times. The Expedata software system allows for macro functions which will produce standardized output formats. Macro 13 provides users with the metabolic variables of interest at each reading for each cage. *CalR* can import data generated by Macro-13 processing.

#### TSE’s LabMaster

LabMaster produces an output which is formatted as normalized to total body weight, e.g., VO2(1), allometric scaling to approximate normalization to lean body mass VO2(2), or uncorrected values VO2(3). *CalR* uses this latter set, the uncorrected values for VO_2_(3), VCO_2_(3), and EE, H(3). To select the variables suitable for *CalR*, go to the “View” menu and select the following parameters: XT+YT, XA, YA, H(3), VO_2_(3), VCO_2_(3), RER, Drink, Feed, and Weight. Also, make sure that the “Export table” setting is “Format 1”. When ready to export, enter the “Export” menu and navigate to **Export > Table** and set “Save as type” to be “.CSV”.

### Statistical approach

*CalR* implements the general linear model to describe the effect of mass on energy expenditure. Experiments in which groups have similar body mass (or body composition), the difference in EE between groups can be analyzed by a one-way ANOVA GLM (Figure 1A). When body mass is significantly different between groups, these masses are included as a covariate, as is commonly recommended for studies of obesity [5, 9–18]. However, one essential requirement for ANCOVA is no difference in *interaction* between EE and mass, i.e. the slopes of the groups must be parallel (Figure 1B). How do we interpret an experiment in which the slopes are not parallel, groups have different interactions between EE and mass, and the assumptions of ANCOVA have been violated? The GLM using a dependent covariate (Type III ANOVA) can adequately account for this scenario

**Figure 1.**
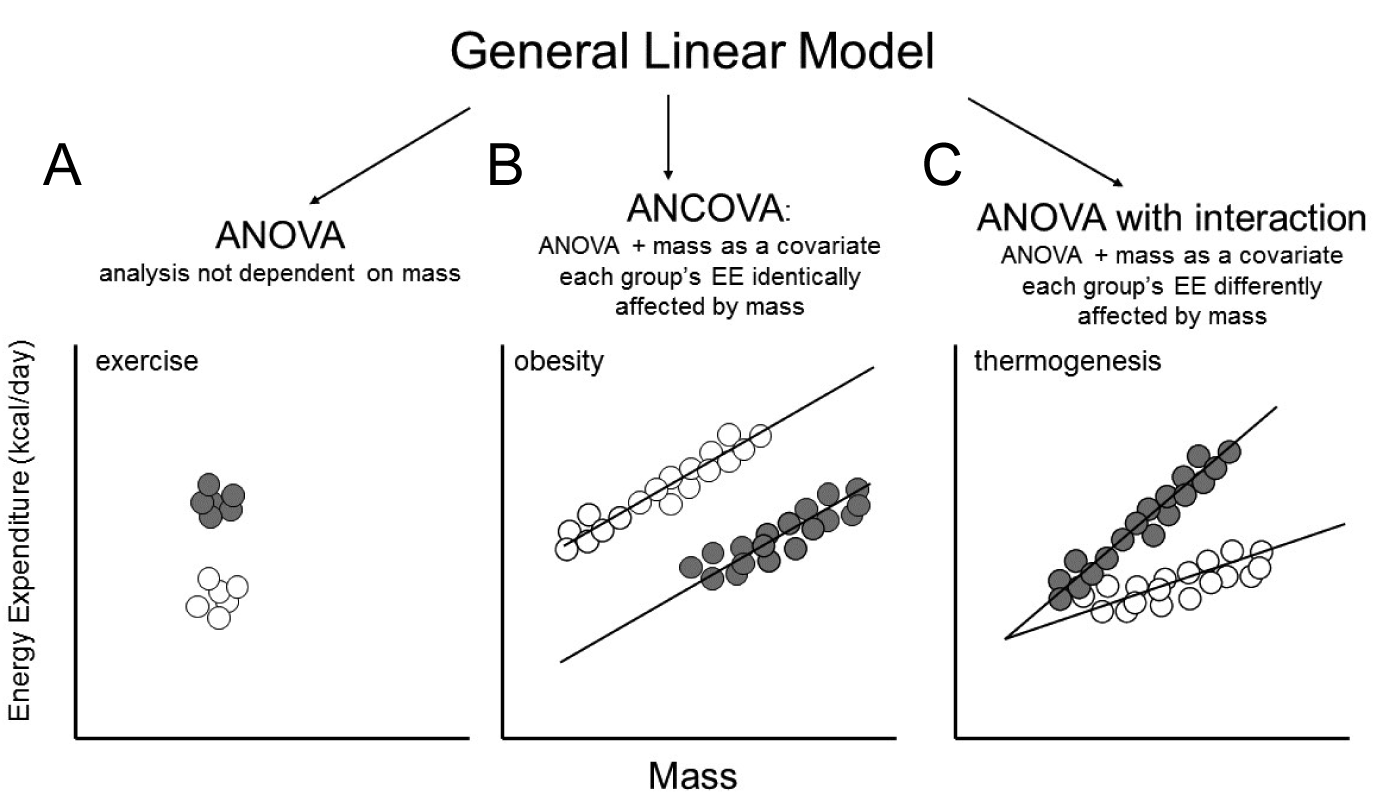
*Analysis models of energy expenditure based on the general linear model*. *CalR determines the appropriate statistical model from the experimental data. A) ANOVA is applied where no difference in mass or body composition exists between groups. An exercised group (gray) with similar mass would use the ANOVA. B) The ANCOVA (ANOVA with the addition of a covariate) when mass is significantly different but slopes are parallel. An obese group of mice (gray) with greater mass but lower EE could be interpreted by this model. C) The ANOVA with interaction is capable of examining the different effect of mass on EE between groups. Mice with activation of BAT (gray) could be interpreted by this model*.

(Figure 1C). *CalR* will first determine whether differences in mass exist, then perform an ANCOVA. If a significant interaction effect is observed, then a GLM is performed, and the significance of the group, mass, and interaction effect are all reported. If there is no significant interaction effect, CalR renders an ANCOVA with group and mass effects only. This algorithm will prove applicable to the majority of indirect calorimetry experiments.

Cells with *, **, or *** denote p-values of <0.05, <0.01, or < 0.001 respectively. The GLM performed in R is generalizable to:

~~~
➢ glm(y ~ mass + group, family=gaussian(link=”identity”))
~~~

Where *y* is a metabolic variable in the set of VO_2_, VCO_2_, EE, food or water intake. The user selects mass to be total body mass, lean body mass, or fat mass. A type III model first computes the sum of squares, but if the interaction effect of group and mass is found to be insignificant, then it is dropped from the model. The two assumptions of this analysis are i) for a Gaussian variability distribution of measurement data and ii) an “identity” link function between the predictor variables and expected values of the metabolic response variable to resemble a classical linear model without transformation.

#### ANOVA

The Analysis of Variance (ANOVA) is performed on parameters measured which are not strictly linked to body mass. The GLM model is reduced to an ANOVA with ‘group’ as the sole predictor variable. From the group names listed in the "Input" tab, the first will be the reference group when this categorical variable is coded into the model.

The ANOVA performed in R is generalizable to:

~~~
➢ glm(z ~ group, family=gaussian(link=”identity”))
~~~

Where z is a metabolic variable in the set of respiratory exchange ratio (RER), locomotor activity, ambulatory activity, body temperature, or wheel running. These variables are independent of mass.

#### Post hoc

For experiments with analysis of more than two groups, Tukey’s honest significant difference post hoc test is performed and graphed to display confidence intervals. For the selected metabolic variable, the “Analysis” tab presents the mean difference with a 95% confidence interval for all pairwise group comparisons.

~~~
➢ glm.model <- glm(y ~ mass + group)
➢ glht(glm.model, mcp(group="Tukey"))
~~~

#### Automatic outlier detection

Within the time range selected, the group means and standard deviations are calculated, stratified into the light and dark photoperiods. If “Yes” is selected for the “remove outlier” radio button, the values that fall beyond three standard deviations from the group mean for the respective light/dark period will be excluded. Since VO2, VCO2, EE, and RER are interdependent, then the removal of any data for one of these variables will lead to the removal of the data for all of them at the corresponding time point.

#### Manual cage exclusion

Users are given the option to exclude any cage starting from a designated hour manually. This feature is designed for use when mice must be removed from a cage that remains actively measured and to prevent data from empty cages being included in the analysis. The “Subject Exclusion” tab is automatically populated with all subject names from the raw data, and with the hour in the recorded data at which the experiment concludes (i.e. no data excluded). The input field should be updated for the subject to contain the hour at which the omission of their measurements should begin. While data from excluded cages will be omitted from analysis, no data are removed or excluded from CalR data files. All excluded data points are saved in an exportable data file and combined with automatic outliers, when in use.

#### Metabolic variables vs. time

The data read into *CalR* is cropped to the hour range selected by the user. To generate hourly time plots, *CalR* subsets the data by either group or subject, depending on user input, and computes averages and standard errors at each hour for the metabolic variable being plotted. The values for the daily bar plots are calculated for each group at each photoperiod (light, dark, or full day) and stratified by day, while the overall bar does not stratify by day. The structure of the data used for analysis consists of the average value of the metabolic variables for each subject for any one of the photoperiods.

#### Body masses and compositions

*CalR* will automatically conduct unpaired two-sample t-tests on all two-group combinations to compute p-values that indicate if there are any differences in the body mass averages of the groups.

~~~
➢ t.test(mass ~ group)
~~~

If in addition to total body mass, the lean and fat masses are included, then *CalR* will also conduct similar statistical analyses of the body compositions. Bar plots display the average masses (and composition), and the number of stars above the bars represents statistical significance. When more than two groups are involved, the significance is pairwise and the comparisons being made are indicated by the start and end location of the horizontal lines above the bars.

#### Time plots

The hourly trends are represented by a line plot shown for each group or subject, depending on the viewing option selected by the user. The daily means are shown directly below the hourly trends and are stratified by the light/dark cycle, displaying the group averages and standard errors by bar plots with error bars. This daily trend plot is replaced by the hourly changes plot in the “two groups, acute response” template. The hourly changes plot can be set to display the absolute or percent difference from the initial value.

#### Regression plot

*CalR* plots the average EE against the mass variable (lean, fat, or total) for the subjects included in the data. The default variable shown is EE because it is less prone to contain error than VO2 and VCO2 in open circuit indirect calorimeters [14]. However, *CalR* provides the option to plot any of the following metabolic variables against body mass: VO^2^, VCO^2^, EE, RER, cumulative food intake, locomotor activity, ambulatory activity, and body temperature [21]. To examine the association between the metabolic variable and mass, we generate a plot with the average of each subject’s mass against the average value of the selected metabolic variable of the respective subject over the experimental time frame chosen by the user. Lines of best fit are produced for each group, from which the slopes are computed and compared by linear regression analysis. For the selected metabolic variable, time of day, and mass variable, *CalR* will automatically calculate the p-values of GLM-based coefficients to indicate if, between the groups, the regression lines are significantly shifted or are not parallel.

## Results

### Typical workflow

The information required to plot and analyze an experiment includes the raw calorimetry data file, body weight or body composition information, and to which group each subject belongs. *CalR* accepts raw indirect calorimetry data from the following manufacturers: Columbus Instruments’ CLAMS, Sable Systems’ Prometheon, or TSE’s LabMaster. The raw data files are parsed and read into memory. The next step is optional as the user chooses whether to import supplemental weight values (i.e. updated body weights). If yes, an option is provided to include either total mass alone or total mass and body composition values. A spreadsheet pre-formatted with animal identifiers is downloaded, filled in with the corresponding mass data, and uploaded. Lastly, group names are specified, the animal identifiers are listed and moved into the corresponding column (Figure 2A). Users can then explore the data under the “Time Plots” tab. Further preferences are subject to users’ selections of inputs including which metabolic variable, plotting by group or individual, the time range, inclusion of error bars (+/-SEM), removal of outliers, and aesthetic features. The raw data is reformatted into a *CalR* raw data file, and all aesthetic features are included in a *CalR* session file. These files obviate the need for repeated data entry in subsequent sessions (Figure 2B). Once these are specified, the tabs producing analyses, including weight plots, regression plots, and GLM results are populated.

**Figure 2.**
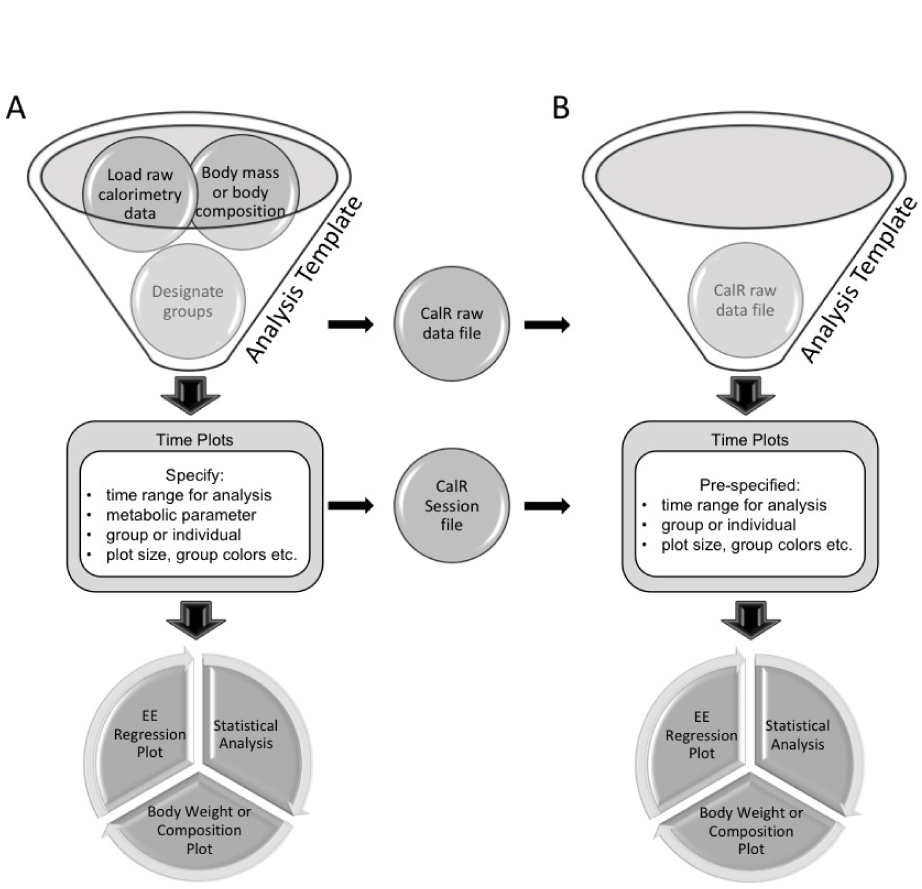
*CalR* data analysis work flow. Users will select an analysis template which best matches the experimental design. A) First time analysis of an experiment includes loading raw indirect calorimetry data, loading body composition data, and allocating animals into groups. This raw data can be exported in a standardized *CalR* data file for fast loading in subsequent sessions. Plotting parameters including the time range for analysis and other aesthetic preferences are set and visualized. These settings are saved into a *CalR* Session file. Once parameters are defined, statistical analysis and additional plotting results are available. B) Exported *CalR* raw data files and *CalR* Session files allow fast, reproducible analysis and for deposition in a data repository.

### Defining an experiment

Each indirect calorimetry run may contain more than one experimental intervention. We present an example in which mice are maintained at thermoneutrality (30˚C) for three days followed by a transition to 4˚C, (Figure 3). The time corresponding to the experimental period of interest can be selected with a slider bar under the “Time Plots” tab (see vignettes). In providing a generalized framework for analysis, this single experiment analysis offers the greatest flexibility for a range of experimental designs.

**Figure 3.**
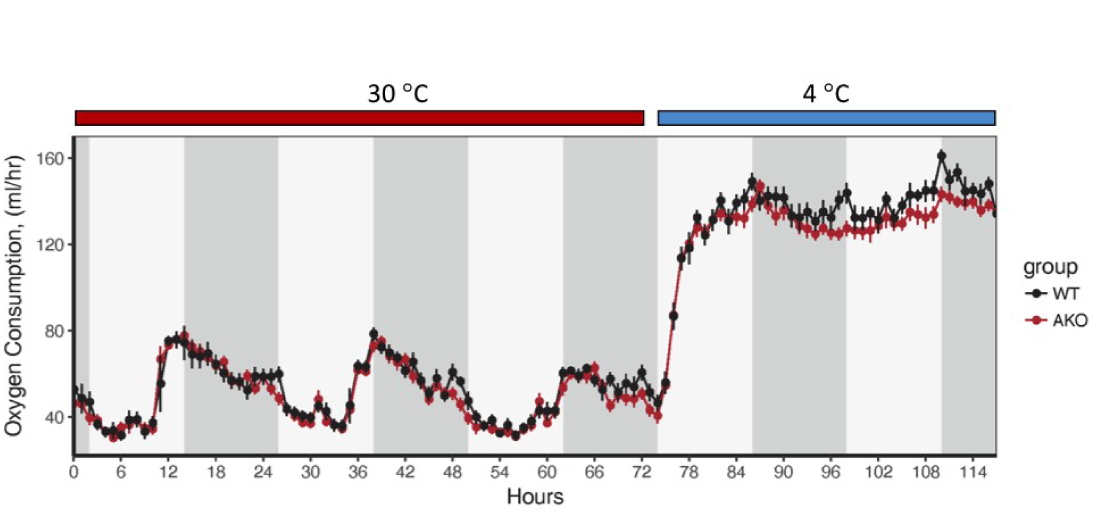
Defining an experiment. This calorimetry run included two experiments in which two groups of mice are maintained at thermoneutrality (30°C) (Experiment 1) followed by a transition to a cold challenge and maintenance at 4°C (Experiment 2). Users of *CalR* will sequentially analyze these two experiments by selecting the corresponding time regions.

### Analysis and tool templates

*CalR* provides the flexibility to interpret many common experimental designs. Each indirect calorimetry experiment is unique, but we have created standardized templates for many common practices (Figure 4). Users are directed to select a template which will perform the most suitable statistical approach. For this reason, we have prepared templates to analyze the following five commonly encountered experimental designs. A description of each template including a sample data set and step-by-step instructions are included (Supplemental File 1):

**Figure 4.**
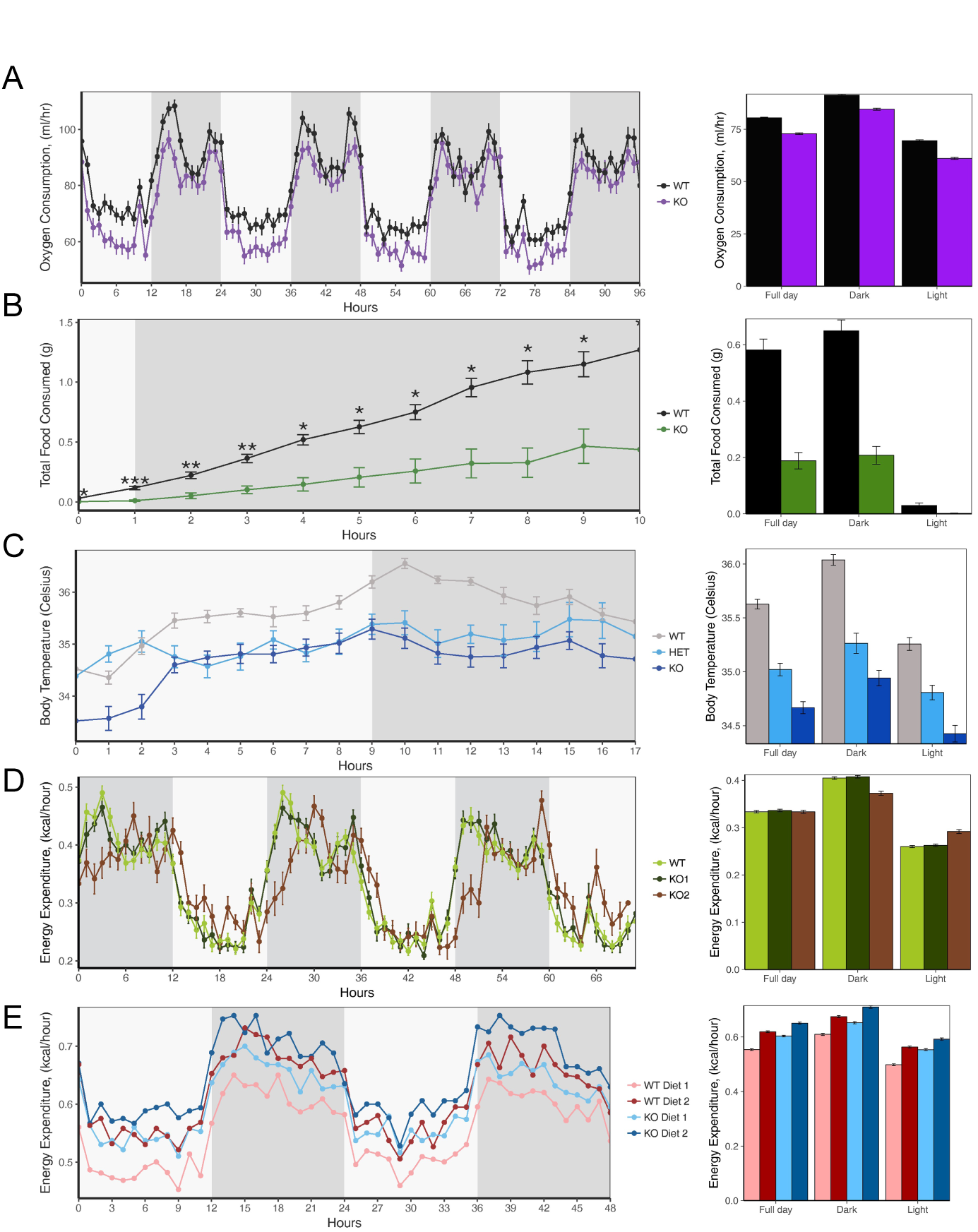
Example data from each of the 5 analysis templates. Left, Time Plot; Right Overall Summary. A) Two group template. VO2 for two groups of mice monitored for four days at room temperature. B) Two group acute response template. Food intake for two groups over 10 hours following treatment. C) Three group template: ordered. Body temperature in wildtype, heterozygote, and knockout animals maintained at 4°C. D) Three group template: non-ordered. Energy expenditure of wildtype and two independent knockout strains. E) Four group template. Energy expenditure analysis of two genotypes of mice on two different diets. Note: *CalR* plots are not normalized or adjusted to body weight, lean mass, or other allometric scaling.

1. **Two groups of mice** (e.g., two genotypes). This is the most common paradigm during which two groups are studied simultaneously.
2. **Two groups with acute treatment** (e.g., administration of a beta-adrenergic receptor agonist to stimulate metabolism). This template is ideal for targeting analysis over a region of 12 hours or less. It includes time plots of metabolic differences from a designated start hour.
3. **Three ordered groups** (e.g., dose-response or wildtype/heterozygous/knockout). This template is for observing dose effects, either allelic, pharmacologic or conditioning. Groups are ordered into a hierarchy for analysis, which includes post hoc tests.
4. **Three factored group** (e.g., Vehicle vs. two independent treatments or WT vs. two independent KOs). Contrary to the ordered template, this one does not assume a hierarchical ordering of the group variable. This makes the analysis, including post hoc tests, distinct from the previous template.
5. **Four groups** (e.g., two genotypes with two diets, or four independent genotypes). This template allows for any combination of four independent comparators and includes post hoc tests.
6. **Experimental run combination Tool.** *CalR* also provides a template which generates a graphical interface to facilitate combining multiple experimental runs into one *CalR* data file.

### Exclusion of recorded data

*CalR* contains features to control automatic and manual exclusion of data. Data specified as excluded will no longer be plotted, calculated into group averages or statistical analysis. However, data is never removed or lost from the *CalR* file. Furthermore, the manually and automatically excluded data will be included in an “excluded data” file.

#### Manual subject exclusion

After beginning a calorimetry run, animals may require veterinary intervention, or have reached a humane endpoint for exclusion or euthanasia according to institutional guidelines. When one animal is removed from a cage, the data collected from this empty cage should cease to be included in the group analysis. Also, the indirect calorimetry apparatus is both complex and error-prone. Exclusion of data from malfunctioning feeders (e.g., readings of negative food intake) is justified, as is the exclusion of data from improperly sealed chambers. Manual data exclusion is designed to remove all data from an empty cage at a designated time for the duration of the experiment.

#### Automatic data exclusion

If the “Exclude Outliers” button is activated, values of more than three standard deviations of the group mean over the time selected will not be included in plots or analyses (see Methods).

#### Data visualization

There are several tabs available in the *CalR* web pages for observing the data with distinct perspectives. Navigating the tabs, a user can see body weight bar plots, time plots, and regression plots. The volumetric data presented are not normalized by total body mass, lean body mass or any allometric scaling factor due to the significant distortions that these can introduce. The values presented under the “Time Plots” tab are the mean values for each group per hour. Also, the mean value for each day and 12-hour light or dark periods are presented as well as overall means for the selected region of analysis. The experimental time period of interest can be specified with a slider bar to perform analysis on one experiment at a time. Many of the features of the time plots can be customized using the dialog box to specify colors, sizes, or inclusion of error bars. Weight plots represent the mean body mass or body composition, if available. Regression plots are often informative for understanding the relationship between EE and mass.

### Transparency, Portability, and Reproducibility

#### The *CalR* Data file

The data files produced by indirect calorimeters of different manufacturers are formatted differently. However, data loaded into *CalR* from each of the three manufacturers can be exported in a standardized format, a *CalR* data file. The *CalR* data file contains the raw data for each animal with hourly averages. The decreased complexity of this file produces a faster computational performance. As a complete standardized record of the experiment, a *CalR* data file can be shared or included as supplemental data.

#### The *CalR* Session file

Outside of the raw calorimetry data, additional information is often necessary to complete analysis. This includes the body weights or body compositions, group placement, cages being excluded from the analysis, time selection, and aesthetic choices. The session file allows for specific and reproducible analysis of either raw data or from a *CalR* file. Multiple *CalR* Session files should be produced for calorimetry runs with numerous distinct experiments. We have included examples of generating and reading the *CalR* file in the vignettes included as supplemental file 1. This modular format will considerably facilitate the sharing of information and the creation of repositories of metabolic data sets.

#### Distribution

The *CalR* graphical front-end of this software operates in a browser window and can either be executed by navigating to a web page hosted by the Brigham and Women’s Hospital (metabolic.bwh.harvard.edu) or downloaded from GitHub, compiled and run locally. This code is free to academic users and is provided under a permissive MIT license.

## Discussion

*CalR* provides users with much-needed comprehensive data analysis tools for indirect calorimetry experiments. Using *CalR* will enable easy access to analysis of metabolic cage data using the GLM which should reduce the “recurring problem” where EE is inappropriately normalized by allometric scaling or divided by lean body mass in mice of different body compositions [6, 11].

ANCOVA has been the method of choice for indirect calorimetry experiments as it efficiently models the effect of mass on multiple metabolic variables. However, by definition, the ANCOVA cannot analyze a differential interaction between mass and group on EE (Figure 1). One approach to circumvent these limitations is using the Johnson-Neyman procedure [22] to find and analyze regions where there is no significant interaction between groups. While the Johnson-Neyman procedure permits the ANCOVA to be performed, it is accompanied by the dual drawbacks of i) excluding data and decreasing already limited statistical power and ii) it may compromise interpretations in cases where the interaction effect may be biologically relevant. Although ANCOVA has been widely recognized as a suitable model for indirect calorimetry data analysis, it is important to have the data drive the decision on which models to use. By transitioning from classical linear regression (ANOVA or ANCOVA) to GLM, assumptions of normality and constancy of variance are no longer required [23]. In the GLMs, interaction effects are included when they are statistically significant. Since the interaction effect could be an essential component of an experimental metabolic story, it is added in *CalR* to provide a more complete and applicable analysis.

Furthermore, the shift to a GLM will continue to support the ability of *CalR* to become a standardized and generalizable tool. Currently, the GLM implemented resembles the consensus linear regression model, however, by design the GLM is flexible and can be adjusted to fit the data better. This feature is important as *CalR* becomes publically available and the scope of experiments being analyzed expands. This is a self-sustainable benefit because as more investigators use *CalR*, the better the community can understand indirect calorimetry data and formulate models that describe its variation; it is necessary to consider experiments run at different locations and points in time and with distinct calorimeters.

There are notable caveats to consider with implementing *CalR* for analysis. Good experimental design is critical for reproducible analysis. Guidelines for experimental design for indirect calorimetry are nicely outlined in Tschop et al. [9].*CalR* cannot detect if a calorimeter is out of calibration and may, therefore, return results which are predicated on faulty data. As with any experimental system, quality control is dependent on the rigorous upkeep of the instrument and vigilance of the operator. Even under optimal conditions, animals may become sick, or equipment failure can spoil the appearance of an experiment. *CalR* provides tools which will allow for the exclusion of data from cages where animals have been removed from an experiment for humane reasons. *CalR* also provides the ability to combine multiple experimental runs to help overcome the difficulties in generating sufficient numbers of sex-matched littermates for the study of genetically modified mouse lines. The investigator is responsible for justifying the appropriateness of two groups being joined for analysis, as *CalR* cannot.

*CalR* will allow for sharing of raw data files of experiments between calorimetry platforms. This will enable whole body physiology to join the broader trend in biomedical research led by genomics and transcriptomics. The ability to efficiently share files as supplemental data will foster increased transparency and reproducibility with the cooperation of interested investigators. We propose a centralized repository of *CalR* indirect calorimetry data files would accelerate global research into metabolism and whole body physiology.

Despite the many features of *CalR*, there is still much room for further innovation. Specifically, newer calorimetry systems provide more frequent sampling times and higher-resolution understanding of metabolic parameters. However, this nuance is lost in ANOVA/ANCOVA/GLM analyses. Regardless of the high-resolution time data, the consensus approach of ANCOVA-like analysis depends upon a single mean value per metabolic variable per mouse [9, 10, 19]. Future advances may be possible by implementing time series and body composition into a statistical framework for indirect calorimetry which accounts for these rich data sets. *CalR* opens the door for communicative efforts to determine how indirect calorimetry data should ideally be analyzed under different experimental conditions.

With data loaded into the R statistical environment, many additional analyses are possible and readily implementable. As an open-source project, *CalR* will benefit from feedback from the engaged community of investigators who have been working on the problem of appropriate analysis of calorimetry results. We encourage comments to this pre-print.

## Supplemental Files

1. Template walkthrough and vignettes
2. Sample data for Vignette example 1: Two-group analysis with ANCOVA and Vignette example 2: Two-group analysis with interaction using the general linear model.

### Acknowledgements

Testing of this program was supported by help from Louise Lantier (Vanderbilt), Dimitrije Cabarkapa (BWH), Barbara Calderone (BWH), C.J. Bare (Cornell), Terry Maratos-Flier (BIDMC), Marie Mather (BIDMC), Bhavna Desai (BIDMC), Jason Kim (UMass), Hye Lim Moh (UMass), Tom Balon (BU), Joe Brancale (MGH), Maria Agostina Santoro (MGH), Lee Kaplan (MGH), Ben Zhou (MGH), Lianfeng Wu (MGH), Alex Soukas (MGH) and Joe Avruch (MGH).

We are grateful for financial support from the NIDDK Mouse Metabolic Phenotyping Centers (MMPC, www.mmpc.org) under the MICROMouse Program, grants DK076169 and DK115255 and from the Harvard Digestive Disease Center, DK034854.

